# Talker discontinuity disrupts attention to speech: Evidence from EEG and pupillometry

**DOI:** 10.1101/2021.01.28.428718

**Authors:** Sung-Joo Lim, Yaminah D. Carter, J. Michelle Njoroge, Barbara G. Shinn-Cunningham, Tyler K. Perrachione

**Author notes:** **Correspondence**, Sung-Joo Lim, PhD, Department of Psychology, Binghamton University, State University of New York, Tyler K. Perrachione, PhD, Department of Speech, Language, and Hearing Sciences Boston University.

## Abstract

Speech is processed less efficiently from discontinuous, mixed talkers than one consistent talker, but little is known about the neural mechanisms for processing talker variability. Here, we measured psychophysiological responses to talker variability using electroencephalography (EEG) and pupillometry while listeners performed a delayed recall of digit span task. Listeners heard and recalled seven-digit sequences with both talker (single- vs. mixed-talker digits) and temporal (0- vs. 500-ms inter-digit intervals) discontinuities. Talker discontinuity reduced serial recall accuracy. Both talker and temporal discontinuities elicited P3a-like neural evoked response, while rapid processing of mixed-talkers’ speech led to increased phasic pupil dilation. Furthermore, mixed-talkers’ speech produced less alpha oscillatory power during working memory maintenance, but not during speech encoding. Overall, these results are consistent with an auditory attention and streaming framework in which talker discontinuity leads to involuntary, stimulus-driven attentional reorientation to novel speech sources, resulting in the processing interference classically associated with talker variability.

## Introduction

There is immense variability in the acoustics of speech across talkers and contexts. The lack of direct, one-to-one correspondences between speech acoustics and linguistic units (Hillenbrand et al., 1995) presents a challenge for listeners who must resolve this inherent variability in order to efficiently perceive and remember speech. Speech processing is less efficient when the source of the speech signal is variable: many prior studies show that listeners are slower and less accurate in processing speech spoken by a series of multiple, mixed talkers compared to processing a single, consistent talker’s speech (e.g., Choi et al., 2018; Mullennix and Pisoni, 1990; Nusbaum and Morin, 1992). This highly replicated phenomenon has been a driving force behind many contemporary psycholinguistic models of speech perception, which attempt to account for how listeners reliably decode speech signals in the face of acoustic variability across talkers (Pierrehumbert, 2003; Kleinschmidt & Jaeger, 2015).

Compared to the behavioral literature, research into how the brain processes speech variability is in its infancy. Functional brain imaging studies have consistently shown greater neural activation in bilateral superior temporal cortex when listening to speech from mixed talkers compared to a single talker (Belin and Zatorre 2003; Wong et al., 2004; Chandrasekaran et al., 2011; Perrachione et al., 2016). These observations support the idea that processing mixed and discontinuous talkers’ speech is less efficient (i.e., there is greater neural activation) than processing a single talker’s speech (Wark et al., 2007; Grill-Spector et al., 2006). Despite the observation of greater superior temporal activation to mixed talker speech, there is no consensus as to why mixed-talker speech incurs additional processing costs. Among the prevailing psycholinguistic models that address the challenge of talker variability in speech processing, each posits a distinct cognitive mechanism to account for the added processing demands, including computational complexity (Nearey, 1989; Johnson, 2005), allocation of cognitive resources (Nusbaum and Magnuson, 1997; Magnuson and Nusbaum, 2007), accessing long-term memory representations (Kleinschmidt and Jaeger, 2015), and stimulus-driven allocation of auditory attention (Bressler et al., 2014; Choi and Perrachione, 2019; Kapadia and Perrachione, 2020). However, prior neuroimaging studies were not designed to test the predictions of different psycholinguistic models. Conversely, the psycholinguistic models were derived principally from behavioral data, and rarely informed by neural evidence or putative neurocomputational mechanisms.

In this paper, we draw upon a range of psychophysiological techniques to evaluate the neural signatures of talker variability with respect to those predicted by a stimulus-driven auditory attention account of talker-specific speech processing (Choi and Perrachione, 2019; Kapadia and Perrachione, 2020). This framework proposes that the increased processing costs for parsing mixed talkers’ speech reflect the cognitive costs involved in switching attention from one auditory source to another (Best et al., 2008; Bressler et al., 2014; Mehraei et al., 2018). Featural discontinuities in the acoustics of an auditory stream—such as those introduced by a switch from one talker to another—disrupt the automatic build-up of a coherent perceptual sound “stream” (Bregman, 1990). Therefore, when listening to mixed talkers’ speech, discontinuities in the source of speech (i.e., talker switches) require listeners to reorient their attention to the acoustic features of each newly encountered talker, which adds processing demands, reflected in lower accuracy and increased reaction time for identifying the linguistic content of speech. Briefly, parsing mixed talkers’ speech interferes with efficient speech processing because talker discontinuity disrupts listeners’ attentional focus, while listening to a coherent speech stream enhances attentional focus (Best et al., 2008; 2018; Bressler et al., 2014; Mehraei et al., 2018).

A growing body of behavioral research supports the view that the processing costs associated with talker variability are better explained by attentional disruption than by phonetic recalibration or increased working- or long term-memory demands. For instance, any discontinuities in talkers—irrespective of the number of talkers or listeners’ expectation about the upcoming talker—interfere equally with speech processing efficiency (Carter et al., 2019; Kapadia and Perrachione, 2020). Furthermore, continuity of acoustic features in a stream of events produces an automatic build-up of attention over time (Shinn-Cunningham, 2006; Sussman et al., 2007; Winkler et al., 2009; Best et al., 2018); however, this build-up depends upon the timing of the discrete events. When events are temporally close, streaming of similar events is enhanced, leading to more efficient speech processing (Best et al., 2008; Bressler et al., 2014; Choi et al., 2018; Lim et al., 2019); conversely, interference from mixed talkers’ speech is greater with temporally contiguous speech sounds, as talker switches become more disruptive when listeners must quickly reorient their attention to each newly encountered talker (Best et al., 2008; Lim et al., 2019). However, behavioral evidence alone does not reveal whether attention-related brain processes are impacted by talker discontinuity, and how such impact changes with temporal proximity of speech.

Beyond its deleterious effects on the immediate recognition of speech, little is known about how talker variability persists in short-term memory for speech information. Recent work has demonstrated that speech working memory retains not only the abstract content of speech, but also other stimulus-specific details (Lim et al., 2015; 2018; 2021), and that working memory for speech is supported by some of the same neural process responsible for speech perception (Jacquemot and Scott, 2006; Perrachione et al., 2017; Scott & Perrachione, 2019). Thus, the impact of acoustic featural continuity across time may help reveal not only how speech is immediately perceived, but also how it is encoded and maintained in working memory. In particular, if both talker and temporal continuity impact listeners’ perception of speech as a coherent stream, discontinuities in these attributes might also interfere with working memory maintenance of speech. While prior behavioral evidence is consistent with this possibility (Martin et al., 1989; Best et al., 2008; Bressler et al., 2013; Lim et al., 2019), disentangling whether discontinuity-related interference arises only during speech encoding, or whether it persists during speech memory retention, is best addressed by physiological, not simply by behavioral, measures.

To evaluate the stimulus-driven attentional streaming hypothesis of processing talker variability, we examined whether the neurophysiological signatures of processing mixed-talker speech are consistent with those associated with disrupting and reorienting listeners’ attentional focus. We also explored whether the added costs of processing mixed-talker speech persisted in the neural signatures of working memory maintenance. To this end, we utilized scalp electroencephalography (EEG) and pupillometry to measure several temporally resolved neurophysiological signals that reflect distinct neural mechanisms for perception and cognition, including cortical evoked potentials, neural oscillatory power, and pupil dilation. We simultaneously recorded EEG and pupillometry while participants performed a delayed recall of digit span task that manipulated two conditions: talker continuity in the speech stream (digits spoken by a single vs. mixed talkers) and temporal continuity in the speech stream (digit streams presented with either 0- vs. 500-ms inter-stimulus intervals; ISIs). Our goals were to (i) assess the impact of talker discontinuity on these neurophysiological signals during task performance, (ii) examine how talker discontinuity-related effects change with temporal discontinuity in speech, and (iii) investigate whether talker discontinuity impacts the maintenance of speech information in working memory.

We hypothesized that several distinct neurophysiological signals would be affected by talker and temporal discontinuities during speech processing and working memory maintenance. First, if processing mixed-talker speech disrupts listeners’ attentional focus and leads to a processing cost associated with reorienting attention to different sources and features, it should elicit the evoked component P3a, which is similar to the distractor positivity (Pd). This is an ERP component that arises around 250–300 ms after an onset of a new stimulus. This component is one of the prominent neural signatures of involuntary attentional reorientation to a distracting (or deviant) stimulus (Comerchero and Polich, 1998, 1999; Polich, 2007; Donchin et al., 1997). Prior work in the visual domain has shown that the P3a/Pd is elicited in response to confusable distractors that share features similar to the target (Hickey et al., 2009; Hilimire et al., 2011; Sawaki and Luck, 2010). Similarly, auditory EEG studies found that this ERP component was elicited when listeners were distracted by a change in task-irrelevant features of a sound (e.g., Goldstein et al., 2002; Gaeta et al., 2003; Stewart et al., 2017). Thus, we expected to observe a P3a-like evoked response to mixed talkers’ speech if listeners’ attentional focus is disrupted by changes in task-irrelevant acoustic features, such as those that occur when the talker switches. Furthermore, if the interference from talker discontinuity depends on the timing of speech, the extent of talker discontinuity-related P3a/Pd response should depend on temporal continuity of the speech stream.

Another signature of attentional reorientation is the phasic pupil dilation response, which indexes the activity of the locus coeruleus norepinephrine (LC–NE) system. The pupil-linked LC-NE system has been associated with arousal and vigilance (Aston-Jones and Cohen, 2005; Sara, 2009; Gilzenrat et al., 2010), as well as cognitive effort and working memory load allocated during task performance (Beatty and Lucero-Wagoner, 2000; Unsworth and Robinson 2015; Kahneman and Beatty, 1966; Heitz et al., 2008; Johnson,1971; Johnson et al., 2014; Peavler,1974). Recent studies also suggest that phasic LC-NE activity plays a role in interrupting an ongoing attentional process to support monitoring of the surrounding environment (Bouret and Sara, 2005; Dayan and Yu, 2006; Sara and Bouret, 2012). Through this mechanism, the pupil-linked LC-NE system is both sensitive to the saliency and surprisal of interrupting sounds (Bala and Takahashi, 2000; Huang and Elhilali, 2017; Liao et al., 2016a, 2016b), and it underlies automatic switching of attention due to an abrupt, bottom-up change in the auditory environment, irrespective of its behavioral relevance (Zhao et al., 2019). Consequently, we expected that pupil dilation responses would reflect the degree to which attentional focus is disrupted and reoriented by talker discontinuity in speech, as well as how the timing of the speech stream affects the degree of disruption.

Finally, to investigate how talker discontinuity impacts working memory maintenance of speech, we also examined how neural oscillatory power, specifically in the alpha frequency range (8–12 Hz), is affected by talker and temporal discontinuity in speech. Alpha oscillatory power has been shown to be a reliable index of the cognitive demands during task performance across various modalities (Jensen and Mazaheri, 2010; Wöstmann et al., 2015; 2016). Specifically, alpha power increases both as more items are held in working memory (Jensen et al., 2002; Tuladhar et al., 2007; Obleser et al., 2012), and when there is an increase in the need to functionally inhibit task-irrelevant processes (Thut et al., 2006; Klimesch et al., 2007; Wöstmann et al., 2015). Thus, if mixed-talker speech imposes greater demand on working memory storage (for instance, if mixed-talker speech is maintained as multiple, discrete speech objects, while single-talker speech is stored as a single object), we expect to observe enhanced alpha oscillatory power for maintaining mixed-relative to single-talker speech in memory. Furthermore, if temporal continuity is necessary for the single-talker speech to be stored as a unitary object in working memory, we expect to see greater alpha oscillatory power for the single-talker stimuli with the 500-ms ISI compared to the 0-ms ISI. However, given that talker discontinuity alone is expected to cause digits to be heard as separate events and thus stored individually, this effect of ISI is likely to be absent or reduced for the mixed-talker conditions. Alternatively, we may observe lower alpha power for maintaining mixed- vs. single-talker speech in memory if mixed-talker speech impairs allocation of attention directed to working memory—for instance, because talker discontinuity leads to inefficient storage of information in memory. Similarly, if temporal discontinuity disrupts efficient attentional allocation, we expect to observe lower alpha power for retaining single-talker speech with the 500-s ISI vs. 0-ms ISI in memory.

## Methods

### Participants

Native English speaking adults (*N* = 24; 16 female, 8 male; mean age: 21 years; age range: 18– 30) participated in the study. Participants were recruited through the Boston University online job advertisement system. All participants had normal hearing (≤ 20dB HL along 250–4000 Hz based on in-lab audiometric tests within 6 months) and reported normal vision. Participants provided written informed consent and were paid $15/hour for their participation. All experimental procedures were approved and overseen by the Boston University Institutional Review Board. One participant’s pupillometry data were excluded due to faulty data recording, yielding *n* = 23 for all pupil dilation analyses.

### Stimuli

The stimuli were natural productions of the digits 1–9, recorded from eight native speakers of American English (4 female; 4 male). Each recording was resynthesized to be 550 ms in duration via the pitch-synchronous overlap and add algorithm (PSOLA; Moulines and Charpentier, 1990) implemented in Praat in order to minimize temporal asynchrony in digit sequence presentations. All recordings were normalized to have equivalent root-mean-square amplitude of 65 dB SPL. On each trial, seven randomly selected digit recordings were concatenated to construct a digit sequence appropriate for the task condition. Digit sequences were constrained such that any digit could appear in any position of a sequence, but the same digit could not repeat in adjacent positions within a sequence.

### Task design and procedure

**Figure 1** illustrates the delayed recall of digit span task, based on a previous behavioral study (Lim et al., 2019). This task manipulated the conditions of *talker discontinuity* (mixed-talker vs. single-talker sequences) and *temporal discontinuity* (500-ms vs. 0-ms ISIs) during digit sequence presentation. On each trial, participants heard a sequence of seven spoken digits and then, after a 5-s “hold” period, recalled the sequence in the order of presentation. The digit sequence was either spoken by one consistent talker (single-talker condition) or each of the seven digits was spoken by a different, randomly selected talker (mixed-talkers condition), so that there was no talker repetition within a given sequence. After the 5-s hold period, participants were prompted to recall the sequence, using a computer mouse to click on the digits (in order) on a number pad GUI that appeared on the screen. Participants were instructed to internally rehearse the digit sequence during the 5-s hold period but to refrain from speaking out loud. Throughout the digit encoding and hold periods, listeners were asked to fixate on a black center dot displayed on the computer screen to minimize artifactual ocular movement. In order to reduce artifacts in the EEG and pupillometry data, participants were also verbally instructed to withhold eye blinks as much as possible until the number pad appeared on the screen.

**Figure 1.**
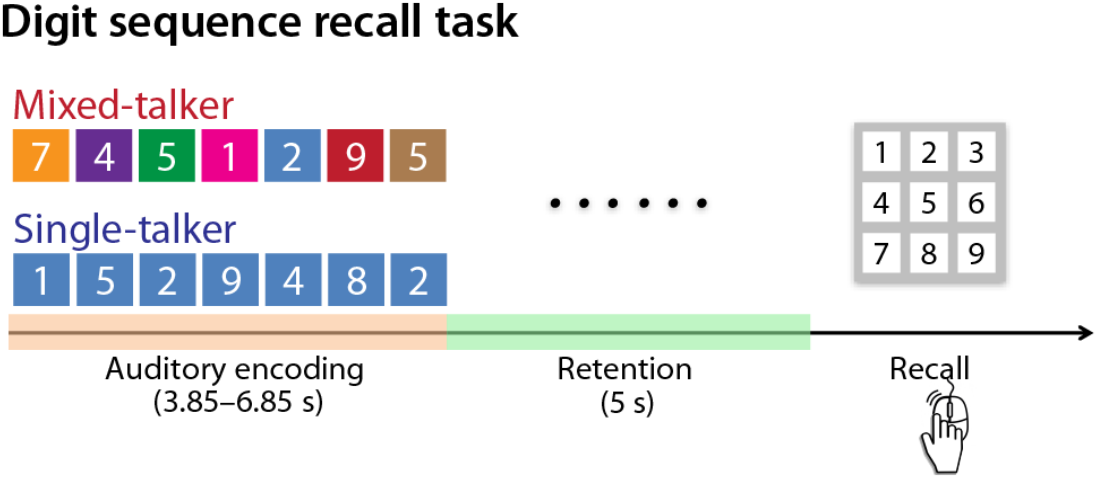
Schematic illustration of the delayed digit span recall task. Listeners heard a sequence of seven digits, either spoken by one consistent talker (*single-talker*) or with each digit in the sequence spoken by a different talker (*mixed-talker*). The digits were presented either with 0-ms or 500-ms inter-digit intervals. After encoding, participants briefly retained the sequence, during a 5-s silent delay period. The appearance of a number pad display on the screen was participants’ cue to begin recalling the digit sequence. Throughout the encoding and retention periods, a center fixation dot was displayed on the screen.

Each participant performed a total of 144 trials. Trials were organized in six runs of 24 trials. Temporal discontinuity conditions (500-ms vs. 0-ms ISIs) varied across runs; all trials in a run had an identical ISI, and the ISI alternated between runs. The order of runs was counterbalanced across participants using Latin-square permutation. Within each run, trials were blocked by talker condition, with three trials per block (i.e., 4 blocks of single-talker and 4 blocks of multi-talker per run). The order of blocks were randomized independently within each run and across participants. We ensured that all speech tokens of each talker were presented an equal number of times across the two talker conditions.

Prior to the experiment, participants completed three practice trials to become familiar with the task structure and with the eye tracker set up. Prior to start of each run, the participant’s head position was stabilized using a head-chin rest, and their eye gaze position was calibrated using a five-point calibration controlled by EyeLink 1000 (SR Research). Participants were given a self-paced break after each trial, and a trial started only after the participant’s eye gaze was stable at the fixation position on the screen; if the fixation drift check failed, eye gaze was re-calibrated before the experiment began. Each trial started 2 s after the eye gaze fixation was accepted, which was indicated to the participants via the color of the fixation dot displayed on the screen.

The experiment was controlled via Psychtoolbox-3 (MATLAB). All sound stimuli were delivered through Etymotic ER-1 insert earphones in a darkened, electrically shielded, sound attenuated chamber. All visual information was displayed via a BenQ 1080p LED monitor (27” diagonal) placed 57 cm from the participant’s head position. The EyeLink 1000 eye tracking system was placed below the computer monitor.

### Data analysis

#### Behavioral analysis

A prior study using an identical task examined listeners’ recall accuracy, response times, and a composite measure of recall efficiency (Lim et al., 2019). The current study focused on only the accuracy measure because task demands of simultaneous pupillometry and EEG recording required participants to suppress blinking until the end of the retention period, precluding them from providing their responses as quickly as possible. Here, we analyzed the accuracy of participants’ memory recall for the digit sequence on each trial. Accuracy was quantified based on correct recall of the digit at each position in the sequence. Data were analyzed in R using a logistic mixed-effects model (*lme4* in R v3.3.3), with recall accuracy of each digit in the sequence as the dependent measure. The model’s fixed-effects terms included the categorical factors of *talker discontinuity* (mixed- vs. single-talker) and *temporal discontinuity* (500-ms vs. 0-ms ISI), the continuous covariate *digit position* in the sequence (1–7) (mean-centered and unit scaled), and all two- and three-way interaction terms. Random-effects terms included by-participant intercepts. Deviation (sum) coded contrasts were applied to the categorical factors. The significance of the effects was determined based on Type-III Wald χ^2^ tests (at *p* < 0.05).

#### EEG data acquisition and preprocessing

EEG data were acquired from 64 active scalp channels in the standard 10/20 montage (Biosemi ActiveTwo system) at a sampling rate of 2048 Hz. Four additional electrodes were placed to record horizontal and vertical ocular eye movements, and two electrodes were placed on the earlobes for reference. Data were downsampled at 1024 Hz and band-pass filtered from 1 to 60 Hz using a zero-phase finite impulse response (FIR) filter (Kaiser windowed, Kaiser *β* = 5.65, filter length 7420 points).

The data were preprocessed using EEGLAB (Delorme and Makieg, 2004). An independent component analysis was performed on continuous data, and artifact components related to eye movements, heartbeat, or noise were identified and removed from the data. On average, 14.5 ± 6.3 (mean ± SD) of 64 components were removed per participant. Data from bad electrodes (2.1 ± 1.7) from seven participants were reconstructed using a spherical interpolation method implemented in EEGLAB. The continuous data were first divided into epochs of −2 to 13 s relative to the trial onset (i.e., onset of the first digit in the trial’s digit sequence). Epochs were removed if any scalp electrodes showed an activity range greater than 200 μV or had infrequent electrical artifacts based on manual inspection within the data segments from −1 to 10 s and −1 to 13 s relative to trial onset for the 0-ms and 500-ms ISI condition trials, respectively. On average, 4.5% of trials were rejected per participant through this procedure; the proportion of remaining trials did not differ in across the experimental conditions [repeated-measures ANOVA; *talker discontinuity*: *F*_1,23_ = 2.07, *p* = .16, *η^2^_p_* = 0.01; *temporal discontinuity*: *F*_1,23_ = .47, *p* = .50, *η^2^_p_* = .014; *talker × temporal discontinuity*: *F*_1,23_ = .34, *p* = .56, *η^2^_p_* = .003]. Data were analyzed using the FieldTrip toolbox (Oostenveld et al., 2011) and customized MATLAB scripts.

#### Event-related potentials

Prior to quantifying event-related potentials (ERPs), the single-trial EEG data were downsampled to 256 Hz. Because cortical auditory evoked responses are sensitive to the acoustic characteristics of speech sounds (e.g., Näätänen and Picton, 1987; Digeser et al., 2009; Khalighinejad et al., 2017; Trembley et al., 2003), the high acoustic variability across the different digit recordings both within a talker and across the eight talkers can increase irrelevant noise in the evoked responses. In contrast to the evoked response to the first digit of a trial, which was preceded by a long period of silence, the evoked responses to the subsequent digits exhibit variable and non-canonical neural response patterns when time-locked to the onset of the audio recordings (see **Figure 3A** and **Figure S1A**). To mitigate this issue and to reduce potential noise, we therefore computed neural evoked responses time-locked to the voicing onsets of the spoken digits in the sequence (**Figure S1B**). First, we divided the single-trial EEG data into 600-ms epochs corresponding to each sequence digit from −0.1 to 0.5 s relative to the voicing onset of the specific digit. These single-trial EEG epochs were baseline corrected by subtracting the mean amplitude in the time window of −0.1 to 0 s relative to the stimulus voicing onset. On average, the onset of voicing varied by 113 ± 24 ms (mean ± SD) from the start of the stimulus recording.

The ERP for each condition was quantified as the average evoked responses to all spoken digits except the first digit (i.e., evoked responses averaged across digits positioned from 2–7). We excluded the response to the first digit because, unlike the other digits in the sequence, the first digit presentation followed a long silence (i.e., the pre-trial period), and the neural response to the first digit in the sequence is independent of the experimental manipulations of interests (i.e., talker discontinuity or temporal discontinuity), which manifest only upon the presentation of the second digit. Confirming these observations, the temporal profile of evoked responses to the first digit was qualitatively and quantitatively distinct from the that of the rest of the digits (**Figure 3A; Figure S1**); also, evoked responses to the voicing onset of the first digit in the sequence did not differ across the conditions (all *p*s > 0.17 from permutation-based cluster tests).

For statistical analysis of ERPs to spoken digits, we used a cluster-based permutation *t*-test provided by Fieldtrip (Maris and Oostenveld, 2007). We conducted three planned analyses, which tested the main effects of *talker discontinuity* and *temporal discontinuity*, and their interaction during the time window of 0 to 0.5 s relative to the voicing onsets of digits. First, to test the main effects of *talker discontinuity*, subject-wise differences between the mean ERPs in the mixed- and single-talker conditions (aggregated across the two temporal discontinuity conditions) were entered into the cluster-based, one-sample *t*-test against zero. Second, to test the main effect of *temporal discontinuity*, we contrasted the mean ERPs of the 500-ms vs. 0-ms ISI conditions (aggregated across the talker conditions) with cluster-based one-sample *t*-test against zero. To test the *talker × temporal discontinuity* interaction effect, we contrasted the ERPs of the talker condition trials (mixed- vs. single-talker) separately for each ISI; this ERP difference was entered into the paired *t*-test to contrast the effect of talker in the 500-ms vs. 0-ms ISI conditions. For each analysis, a permutation test (1000 Monte Carlo permutations) was performed with a cluster-based control at a type I error level of alpha = 0.05, implemented in Fieldtrip. The test resulted in time–electrode clusters exhibiting significant effects of the corresponding contrasts. For any of the significant clusters, we also examined the magnitude of the evoked response elicited in each condition (2 *talker × 2 temporal discontinuity*) using a one-sample *t*-test against 0.

#### Time–Frequency analysis

EEG data were re-referenced to the average of all EEG electrodes. We obtained time–frequency representations (TFRs) of single-trial EEG data by convolving the data with a Hanning taper (fixed window length), which covered frequencies from 1 to 30 Hz with a resolution of 1 Hz. This procedure was applied to the whole epoch from −2 to 13 s relative to the onset of the trial (i.e., the onset of the first spoken digit) using a time step of 0.1s. Single-trial power estimates were baseline corrected by subtracting the relative change with respect to the average oscillatory power during the pre-stimulus baseline (−0.5 to 0 s) across all conditions.

We focused on how talker discontinuity and temporal discontinuity affect neural oscillatory power during working memory retention after encoding digits. We adopted multi-level statistical analysis used in previous studies (Obleser et al., 2012; Lim et al., 2015; Wöstmann et al., 2017) for conducting three planned analyses, testing the main effects of *talker discontinuity* and *temporal discontinuity* as well as their interaction during the 5-s hold period during which listeners retained information in working memory. Single-subject-level statistical analysis was performed on single-trial data using an independent-samples regression *t*-test with contrast coefficients of −0.5 and 0.5 to test the effect of interest (i.e., *talker discontinuity:* mixed- vs. single-talker trials, *temporal discontinuity:* 500-ms vs. 0-ms ISI trials, respectively) while collapsing the other condition; this resulted in the individual subject’s frequency–electrode beta weights of the corresponding contrast, which were entered into the group-level analysis.

To test the main effects of *talker* and *temporal discontinuity* at the group level, a cluster-based one-sample *t*-test against zero was performed from the frequency range from 3 to 20 Hz across the 5-s hold period. For testing the *talker × temporal discontinuity* interaction, we computed beta weights contrasting the talker conditions (mixed- vs. single-talker) on the first-level analysis separately for the 0-ms and 500-ms ISI condition trials; the resulting beta contrasts of the two speech rate conditions were tested at the group level using a paired-sample *t*-test. As when analyzing ERPs, the group-level analysis was conducted as using a cluster-based permutation test with 1000 random iterations (Maris and Oostenveld, 2007). Each test resulted in time–electrode–frequency clusters exhibiting significant effects of the corresponding conditions on the TFRs throughout the 5-s retention window.

In addition, we also examined the main effect of *talker discontinuity* on the neural oscillatory power across 3–20 Hz during the presentation of the spoken digit sequence, henceforth referred to as the encoding phase of the trial, using the multi-level analysis approach as described above. Due to the differences in the digit sequence durations across the two temporal discontinuity conditions, we contrasted the mixed- vs. single-talker conditions separately for the 0-ms and 500-ms ISI condition trials during this encoding phase (0-ms ISI: 0–3.85s, 500-ms ISI: 0–6.85s relative to trial onset).

#### Pupillometry preprocessing and analysis

Pupil diameters from the left and right eyes were continuously recorded with the EyeLink 1000 (SR Research) at a sampling rate of 500 Hz. The pupillometry data were epoched from −2s to 8.85 s and −2 to 11.85 s relative to the trial onset (i.e., first spoken digit) for the 0-ms and 500-ms ISI trials, respectively. Preprocessing steps included a de-blinking procedure and removal of artifact trials. We used a combination of eye position, velocity, and acceleration filters to automatically detect eye blinks, shown as rapid drops in the pupil size traces. The blink data segment was replaced based on linear interpolation between the pupil sizes before and after each blink using customized MATLAB functions. We rejected any trials in which either more than 50% of the data consisted of blinks or that contained long blinks (duration greater than 500 ms) within the 1-s pre-trial window. Subsequently, trials with excessive noise were manually identified and removed from data analysis. On average, 17.35 (±15.49) out of 144 trials were rejected through this process. The pupil diameter of each trial was quantified as the mean of the pupil tracings of the two eyes. For trials in which the fixation of one eye was unstable during recording, the pupil tracing from the other stably fixated eye was used for the trial (3.65% of trials were a single pupil recording).

Pupil diameter during the whole trial period was baseline corrected by computing the relative change from the average pupil diameter during the pre-stimulus baseline (−0.5 to 0 s) of each trial. It is of note that there were no significant effects of the conditions or their interaction on the average pupil diameters during baseline [repeated-measures ANOVA; *talker discontinuity*: *F*_1,22_ = 0.60, *p* = .45; *temporal discontinuity*: *F*_1,22_ = 1.17, *p* = .29; *talker × temporal discontinuity*: *F*_1,22_ = 1.16, *p* = .29].

Our main question was whether pupil dilation responses during task performance were sensitive to talker and temporal discontinuity during the speech encoding and the memory retention phases of the trials. To assess pupil dilation responses during encoding of spoken digits, we quantified mean pupil diameter during the presentation of each digit in the sequence of each trial. In order to account for the temporal delay in pupillary response from stimulus onset (Hoeks and Levelt, 1993; Verney et al., 2004; Winn et al., 2015; 2018), the time windows in which pupil diameters were averaged extended 500 ms beyond the onset of each digit in the sequence. In order to quantify pupil responses during memory retention, we averaged pupil diameters during the 5-s memory retention phase of each trial.

The resulting trial-wise pupil diameter data were analyzed using linear mixed effects models. The fixed effects structure included categorical factors for *talker discontinuity*, *temporal discontinuity*, and their interaction. An additional continuous fixed factor for *digit position* (1–7; mean-centered and scaled), as well as its interaction with the other fixed factors, was entered into the model for analyzing pupil responses during encoding of spoken digits. The random effects structure included by-participant intercepts. For any significant effects, post-hoc tests were conducted via pairwise differences of least-square means (*difflsmeans* implemented in *lmerTest* in R).

## Results

### Talker discontinuity in speech interferes with speech working memory recall

We analyzed whether the 2 *talker* × 2 *temporal discontinuity* condition manipulations affected performance on the digit sequence recall task. The results of the logistic mixed-effects model of participants’ memory recall accuracy are listed in **Table 1**. The model revealed a significant main effect of *talker* [*χ*^2^(1) = 22.35, *p* ≪ 0.0001], but no significant effects of *temporal discontinuity* [*χ*^2^(1) = 1.46, *p* = 0.23] or the interaction *talker × temporal discontinuity* [*χ*^2^(1) = 0.030, *p* = 0.86]. These results indicate that performance was less accurate for digit sequences spoken by mixed talkers than by a single talker; however, the effect of talker discontinuity on memory recall accuracy did not depend on the ISI of spoken digits (**Figure 2**).

**Figure 2.**
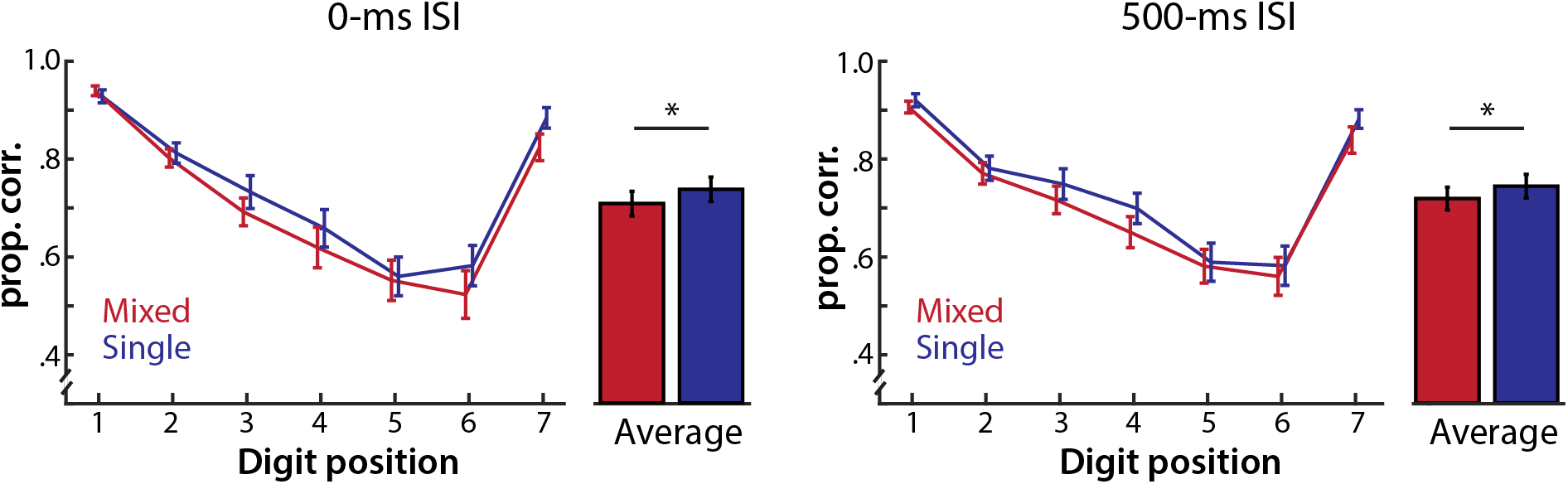
Digit sequence recall performance in the talker and temporal discontinuity conditions. Mean recall accuracy for each digit position is illustrated in the mixed- vs. single-talker conditions in the 0-ms ISI (left) and 500-ms ISI (right) conditions. The bar graphs show mean performance across all digit positions. The error bars indicate ±1 standard error of mean (SEM) across participants. * *p* < 0.05.

**Table 1.** Mixed-effects logistic modeling results of the behavioral recall accuracies across digits in sequences

| Fixed factors | $\chi^2$ | df | p |
| --- | --- | --- | --- |
| Talker discontinuity | 22.35 | 1 | $\ll 0.0001$ |
| Temporal discontinuity | 1.46 | 1 | 0.23 |
| Digit position | 426.51 | 1 | $\ll 0.0001$ |
| Talker × Temporal discontinuity | 0.030 | 1 | 0.86 |
| Talker discontinuity × Digit position | 3.21 | 1 | 0.073 |
| Temporal discontinuity × Digit position | 5.23 | 1 | 0.022 |
| Talker × Temporal discontinuity × Digit position | 1.40 | 1 | 0.24 |

Note: Type III Wald $\chi^2$ test

The model also revealed a significant main effect of *digit position* [*χ*^2^(1) = 426.51, *p* ≪ 0.0001]. As illustrated in **Figure 2**, listeners’ recall accuracy was greatest for recalling digits presented in the initial and final positions of the sequence, exhibiting typical primacy and recency effects of sequence recall. Importantly, the significant interaction between *temporal discontinuity* and *digit position* [*χ*^2^(1) = 5.23, *p* = 0.022] shows that the pattern of recall accuracy across sequences differs for the two ISIs: while participants were significantly more accurate in recalling the first two digits in the sequence with 0-ms ISI compared to with 500-ms ISI [*b*s > 0.099, *zs* > 2.31; *p*s < 0.021], the 500-ms ISI condition led to higher accuracy in recalling the digit in the medial position (i.e., digit position 4: *b* = –0.089; *z* = −2.35, *p* = 0.019).

### Evoked responses during speech encoding are impacted by talker discontinuity and temporal discontinuity

**Figure 3A** illustrates each condition’s evoked response time course throughout the trial. Our main interest in this analysis was whether talker discontinuity in speech affected auditory evoked responses and whether the extent of the response differed across the temporally continuous vs. discontinuous speech stream. The first permutation-based cluster test examined the main effect of the talker condition on the average ERPs across the two ISI conditions. This test revealed one significant cluster exhibiting a mixed-talker > single-talker effect (**Figure 4**), such that the mixed-talker condition exhibited a stronger positive potential compared to the single-talker trials in the time range of 215–348 ms following the voicing onsets of spoken digits (*p* = 0.001). This effect was widely distributed, but most pronounced in the fronto-central electrodes.

**Figure 3.**
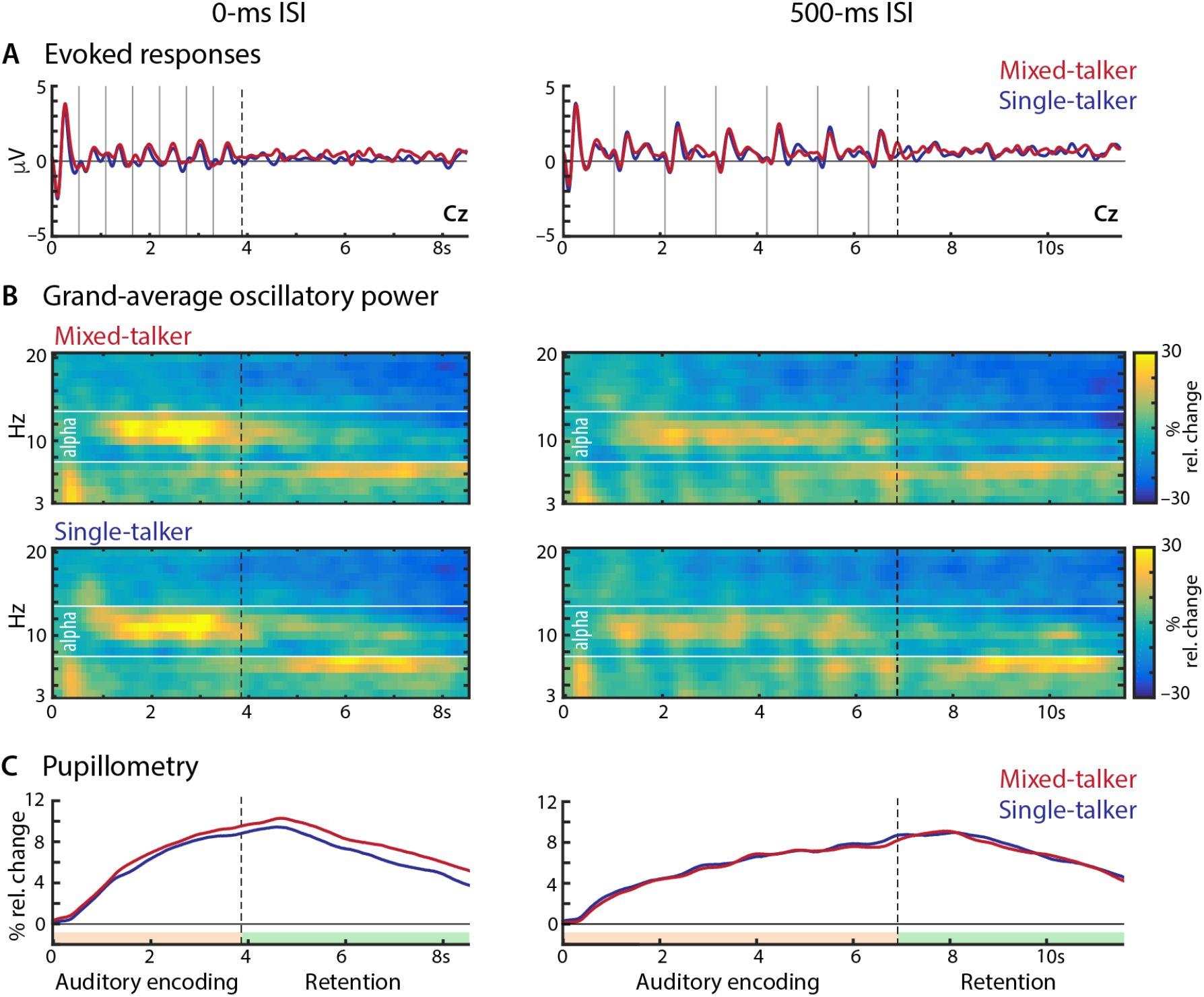
Illustration of the mean time courses of the three neurophysiological responses during the delayed digit sequence recall task. **(A)** Grand average evoked response time courses across participants, time-locked to the onset of the trial extracted from one representative electrode (Cz). For illustration, the grand average time courses were low-pass filtered at 15 Hz. Thin vertical lines denote the onsets of spoken digits in the sequence. Dashed lines mark the offset of the last digit in the sequence (i.e., end of sequence encoding and beginning of retention) as denoted in panel C. Red and blue lines indicate average responses of the corresponding neurophysiological time courses during mixed- and single-talker condition trials, respectively. **(B)** Condition-specific grand average time–frequency representations across participants, and across all scalp channels. The color bar indicates relative oscillatory power change from baseline. Alpha frequency range (8–13 Hz) is demarcated by white horizontal lines. **(C)** Mean pupillometry response time courses across participants in each condition. Y-axis indicate relative pupil dilation change from the average pupil responses during 500ms pre-trial baseline.

**Figure 4.**
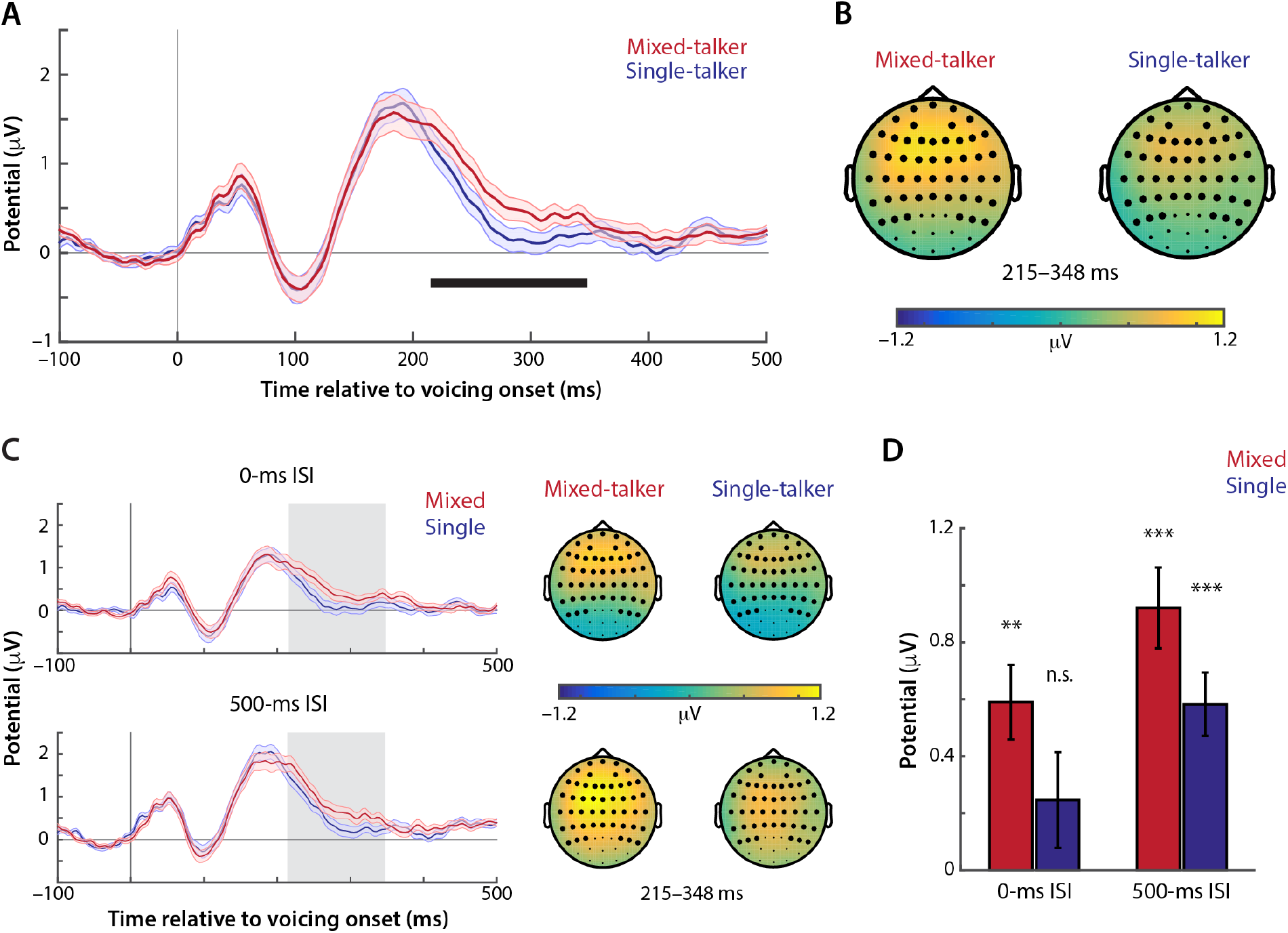
Grand average event-related potentials (ERPs) across participants time-locked to the onsets of voicing of spoken digits in the sequence (excluding the first digit of each sequence). **(A)** Evoked responses in the mixed-talker vs. single-talker trials, averaged across 49 electrodes belonging to the significant cluster. The black horizontal line denotes the time period in which the cluster-level statistic showed a significant main effect of the talker conditions (i.e., significant difference between the mixed- vs. single-talker conditions collapsed across the two ISI conditions). Error bars indicate ±1 SEM. **(B)** Topographical maps of average evoked response amplitudes within the time–electrode cluster exhibiting a significant mixed-talker vs. single-talker contrast. Highlighted scalp channels belong to the significant cluster. **(C)** Grand average time courses (left) and topographical maps (right) of the ERPs in the 2 *talker ×* 2 *temporal discontinuity* conditions. The grey boxes in the time courses denote the same time points that belong to the cluster shown in (A). **(D)** A bar plot of the condition-specific average evoked response amplitudes over electrodes and times of the cluster shown in (A). Comparisons to zero: ** *p*< 0.001, *** *p* ≪ 0.0001.

The other cluster test examining the main effect of temporal discontinuity yielded two significant clusters, both exhibiting stronger positivity in the 500-ms than the 0-ms ISI (**Figure S2**; Cluster #1: 0–63 ms; *p* = 0.015; Cluster #2: 105–500 ms; *p* < 0.001). However, the cluster test evaluating the *talker × temporal discontinuity* interaction effect did not yield any significant clusters. Thus, the pattern of results indicates that for both ISIs, mixed-talker trials exhibited a significantly larger positive potential ~300 ms after the onset of spoken digits than did single-talker trials (**Figure 4C–D**). Based on this pattern, we further examined the magnitude of the response elicited by spoken digits of each condition using a one-sample *t*-test against 0. This test revealed that only the single-talker, 0-ms ISI condition did not yield significantly positive potential (**Figure 4D**; mixed-talker speech at 500-ms ISI: *t*_23_ = 6.49, *p* ≪ 0.0001; mixed-talker speech at 0-ms ISI: *t*_23_ = 4.51, *p* = 0.00016; single-talker speech at 500-ms ISI: *t*_23_ = 5.23, *p* ≪ 0.0001; single-talker speech at 0-ms ISI: *t*_23_ = 1.47, *p* = 0.16).

### Alpha oscillatory power during speech working memory retention is sensitive to talker discontinuity

Our main question in analyzing oscillatory power was whether mixed- and single-talker speech produced different levels of neural oscillations during memory retention, and whether any such effect of talker condition differed across the two temporal discontinuity conditions. **Figure 3B** shows the grand average oscillatory power time courses of the mixed-talker vs. single-talker conditions across the two ISI conditions. In all cases, there is a clear band of power in the alpha range that extends from the encoding phase and into the memory retention phase of the trials. We investigated the main effect of talker discontinuity (mixed- vs. single-talker) during the 5-s memory retention phase using a permutation-based cluster test. This analysis identified one cluster 1.95–3.55s after the onset of the memory retention phase (**Figure 5A**); this cluster exhibited significantly less oscillatory power in the alpha frequency range [8–11 Hz, *p* = 0.001] when listeners were maintaining digits spoken by mixed talkers compared to a single talker in both 0-ms and 500-ms ISI trials (**Figure 5B**). The cluster test examining the *talker × temporal discontinuity* interaction effect did not yield any significant clusters (all *p*s ≥ 0.36). This pattern indicates that compared to maintaining single-talker speech, maintaining mixed-talker speech led to significantly less alpha power during the retention period across both temporal discontinuity conditions.

**Figure 5.**
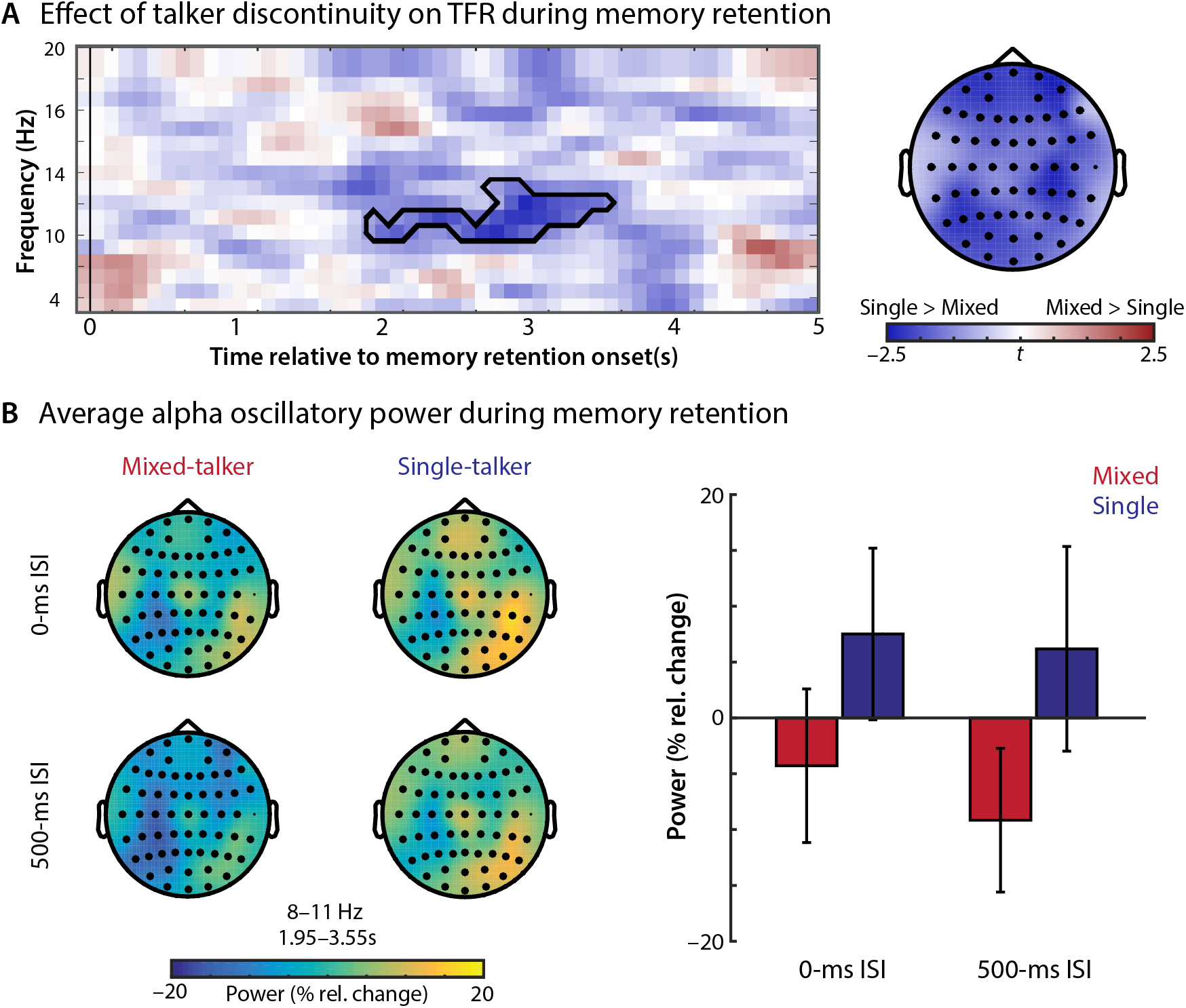
Effect of talker discontinuity on neural oscillatory power during the 5-s memory retention phase. **(A)** Illustrations of the time–frequency representation and topographical map of the cluster exhibiting a significant main effect of the talker condition (i.e., mixed- vs. single-talker contrast aggregated across the two ISIs). The colors indicate the distributions of *t*-statistics of the mixed-talker vs. single-talker contrast. The highlighted electrodes on the topography belong to the corresponding cluster. **(B)** Average oscillatory power in the 2 *talker ×* 2 *temporal discontinuity* conditions over the single-talker > mixed-talker cluster shown in (A). Left, topographical illustrations of the condition-specific averages of oscillatory power over the frequencies and times of the cluster. Right, a bar plot of condition-specific oscillatory power average over the electrodes, frequencies, and times of the cluster. Error bars indicate ±1 SEM.

Although the main effect of *temporal discontinuity* (500-ms vs. 0-ms ISI) was not our main interest, we also noted that the temporal discontinuity of speech across the talker conditions had a significant effect on neural oscillatory power. As shown in **Figure S3**, a permutation-based cluster test examining the main effect of temporal discontinuity revealed a significant enhancement of alpha and lower beta power (10–16 Hz) when listeners were maintaining digit sequences presented at 0-ms ISI compared to 500-ms ISI [0–2.75s; *p* = 0.003] in both talker conditions.

Lastly, we examined whether alpha oscillatory power differed when listeners were encoding speech spoken by mixed-talkers vs. a single-talker, as might be expected under cognitive resource-allocation based models of talker adaptation (Magnuson and Nusbaum, 1997; Magnuson and Nusbaum, 2007). The corresponding permutation-based cluster did not reveal any clusters exhibiting significant differences in neural oscillatory power in the range from 3–20 Hz during speech encoding of mixed- vs. single-talker digit sequences in either *temporal discontinuity* condition trials [0-ms ISI: all *p*s ≥ 0.11; 500-ms ISI: all *p*s ≥ 0.60].

### Pupil dilation response during speech encoding is sensitive to talker and temporal discontinuities

We investigated whether task-evoked pupil dilation responses were related to encoding and maintaining mixed-talker vs. single-talker speech, and whether temporal discontinuity of speech affected pupil responses. **Figure 3C** illustrates the time courses of the pupillary response during task performance in each experimental condition.

We analyzed the effects of *talker discontinuity* and *temporal discontinuity* on the pupillary responses for encoding each digit in the sequence. Linear mixed-effects model (**Table 2**) revealed significant effects of *temporal discontinuity* [*F*_1,20362_ = 24.62, *p* ≪ 0.0001] and *digit position* [*F*_1,20362_ = 1147.95, *p* ≪ 0.0001], as well as their interaction [*temporal discontinuity × digit position*: *F*_1,20361_ = 10.83; *p* = 0.0010]. As illustrated in **Figure 6**, pupil dilations were larger for encoding digits presented at 0-ms ISI than at 500-ms ISI. Also, while pupil dilations generally increased over the course of the encoding phase for both ISIs, the amount that pupil dilation increased per digit was higher in the 0-ms than 500-ms ISI conditions [*temporal discontinuity × digit position: β* = 0.0022].

**Table 2.** Mixed-effects linear modeling results on the pupil dilation responses to spoken digits during the sequence encoding phase

| Fixed factors | *df1* | *df2* | *F* | *p* |
| --- | --- | --- | --- | --- |
| Talker discontinuity | 1 | 20362 | 1.08 | 0.30 |
| Temporal discontinuity | 1 | 20362 | 24.62 | $\ll 0.0001$ |
| Digit position | 1 | 20361 | 1147.95 | $\ll 0.0001$ |
| Talker × Temporal discontinuity | 1 | 20361 | 6.18 | 0.013 |
| Talker discontinuity × Digit position | 1 | 20361 | 0.075 | 0.78 |
| Temporal discontinuity × Digit position | 1 | 20361 | 10.83 | 0.0010 |
| Talker × Temporal discontinuity × Digit position | 1 | 20361 | 0.029 | 0.86 |

*Note:* Type III Analysis of Variance with Satterthwaite's method

**Figure 6.**
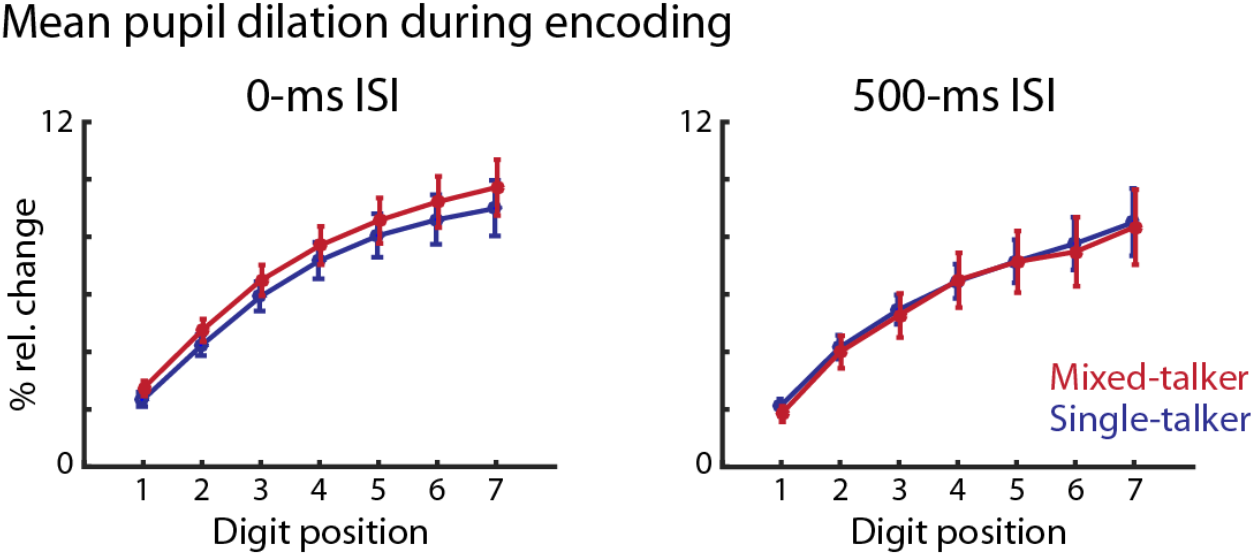
Mean pupil dilations in the 2 *talker ×* 2 *temporal discontinuity* conditions during encoding of each spoken digit in the sequences. The error bars indicate ±1 SEM.

The same linear mixed-effects model did not reveal a significant main effect of *talker discontinuity* [*F*_1,20361_ = 1.08, *p* = 0.30], but there was a significant *talker × temporal discontinuity* interaction effect [*F*_1,20361_= 6.18, *p* = 0.013]. As shown in **Figure 6**, planned post-hoc tests revealed that when digits were presented at 0-ms ISI, pupil dilations were significantly larger for encoding digits spoken by mixed talkers vs. a single talker [*t*_20361_ = 2.52, *p* = 0.012]; however, there was no effect of talker condition on pupil dilation for digits spoken at 500-ms ISI [*t*_20362_ = 1.01, *p* = 0.31].

We also examined how these factors affected pupil dilation during memory retention. We examined the effects of *talker discontinuity*, *temporal discontinuity*, and their interaction on the average pupillary responses during the 5-s memory retention phase. A linear mixed-effects model did not yield any significant main or interaction effects [*talker. F*_1,2887.2_ = 0.86, *p* = 0.35; *temporal discontinuity*: *F*_1,2887.4_ = 0.25, *p* = 0.62; *talker × temporal discontinuity*: *F*_1,2887_ = 1.83, *p* = 0.18].

## Discussion

Although several psycholinguistic models have been put forward to explain how the differences in acoustic-phonetic mappings across talkers place additional processing demands on listeners during speech perception (Nearey, 1989; Johnson, 2005; Nusbaum and Magnuson, 1997; Kleinschmidt and Jaeger, 2015), previous research has not explicitly examined whether the psychophysiological signatures of mixed-talker speech perception are consistent with the distinct cognitive mechanisms proposed by these frameworks (Belin and Zatorre 2003; Wong et al., 2004; Chandrasekaran et al., 2011; Perrachione et al., 2016). In the current study, we examined whether the psychophysiological correlates of encoding and retention of mixed-talker speech were consistent with the cognitive processes posited by one of these frameworks: stimulus-driven auditory attention and streaming (Bregman, 1990; Shinn-Cunningham, 2006; Winkler et al. 2009).

We found that talker discontinuity interfered with listeners’ serial recall of speech from working memory. This behavioral interference from talker discontinuity was accompanied by neurophysiological responses signaling attentional reorientation and interruption of an ongoing auditory streaming process. Encountering discontinuities in incoming speech elicited evoked responses similar to the P3a/Pd ERP component, the magnitude of which increased additively with increased discontinuity from either switches in talkers, temporal separation, or both. Mixed-talkers’ speech led to increased phasic pupil responses, particularly when listeners had to process speech rapidly. Furthermore, compared to single-talker speech, mixed-talker speech produced less alpha oscillatory power during working memory maintenance, but not during initial speech encoding.

Broadly, these results are in line with the neural mechanisms predicted by an auditory attention and streaming framework of talker-specific speech processing (Bressler et al., 2014; Choi and Perrachione, 2019; Kapadia and Perrachione, 2020). Specifically, talker discontinuity interrupts listeners’ attentional focus to an ongoing auditory stream formed by the attended talker; obligatory exogenously driven attentional shifts to novel speech sources encumber the auditory system, resulting in the processing interference classically associated with mixed-talker speech perception.

### Talker discontinuity interferes with speech working memory recall

We found that listeners were less accurate at recalling speech sequences spoken by mixed talkers than by a single consistent talker. This performance interference by talker discontinuity is in line with decades of results demonstrating a consistent and robust interference effect of mixed talkers’ speech on immediate recognition of speech (e.g., Choi et al., 2018; Mullennix and Pisoni, 1990; Nusbaum and Morin, 1992). Behavioral measures alone cannot adjudicate whether less accurate recall of mixed-talker speech is driven solely by inaccurate encoding of mixed-talker speech or also by an influence of talker discontinuity on working memory maintenance (Lim et al., 2019). However, our EEG results, especially our alpha oscillatory power (discussed below), suggest that talker discontinuity continues to impose processing costs even while listeners maintain speech information in working memory (**Figure 5**). Specifically, the fact that there is a significant difference in alpha power when maintaining digits suggests that there are differences in working memory demands for single- vs. mixed-talker stimuli, even after the digits have been encoded and abstracted to memory.

Overall, talker discontinuity did not differentially affect performance accuracy between the temporal discontinuity conditions (**Figure 2**). Superficially, this result seems at odds with those of a prior study from our laboratory: utilizing the identical experimental paradigm, we found that talker discontinuity caused the largest performance disruption for temporally contiguous speech (Lim et al., 2019). The lack of differences across the temporal discontinuity conditions in the present study likely reflects the limited sensitivity of accuracy alone as a dependent variable for describing speech processing dynamics in single- vs. mixed-talker settings. Prior work on immediate speech identification has shown that talker discontinuity has a much greater effect on word identification speed than on accuracy (Kapadia & Perrachione, 2020). Consistent with this, our previous study found that talker discontinuity had a larger effect for temporally continuous speech when considering response time and speech processing efficiency (which simultaneously accounts for the speed and accuracy of memory recall), but not for recall accuracy considered alone. As noted above, we could not reliably assess participants’ memory recall speed in the current study, where we simultaneously measured EEG and pupil dilation. Specifically, in order to obtain artifact-free physiological measures, participants were asked to withhold responses until after the retention period. Moreover, because participants were told to refrain from blinking during the encoding and retention periods, many actually delayed responding until after blinking.

### Talker discontinuity affects neural dynamics of automatic attentional reorientation during speech processing

Across the two temporal discontinuity conditions we found that, compared to a single talker’s speech, listening to mixed talkers’ speech consistently evoked greater fronto-central positive deflections arising about 200–350 ms after the onset of spoken digits (**Figure 4**). This positivity aligns with the P3a/Pd ERP components that are reliably known to appear around ~250 ms after encountering a novel, and potentially distracting, stimulus, as demonstrated in both auditory and visual search tasks (Comerchero and Polich, 1999; Polich, 2007; Sawaki and Luck, 2010). Task-irrelevant novel sounds and/or distractors have been shown to disrupt ongoing task performance following the distractors; the P3a/Pd responses elicited by distractors reflect this involuntary attentional reorientation to a novel stimulus (or a change) in the perceptual environment (Polich, 2007; Escera et al., 1998; Stewart et al., 2017; Sawaki and Luck, 2010). The increased positivity we found during encoding of mixed talkers’ speech suggests that a discontinuity in a task-irrelevant feature (here, talker voice) of a task-relevant stimulus (the digits in the stream) is, on its own, enough to cause a performance-disrupting reorienting response. That is, changes in the task-irrelevant perceptual feature of voice disrupt the encoding of the incoming speech due to an involuntary reorienting to the novel speech source.

We found that both talker discontinuity and temporal discontinuity increased the neural signature of reorienting (the magnitude of the P3a/Pd), but that there was no significant interaction between these two main effects (**Figure 4D**); instead, the effects of talker discontinuity and temporal proximity appear to be additive. It is notable that even when listeners were encoding speech spoken by one consistent talker, temporal discontinuity alone was sufficient to produce a robust P3a response. Temporal continuity of sounds plays an important role in the emergence of auditory streaming (van Noorden, 1975) and temporal gaps (e.g., a 500-ms ISI) between sounds can break down automatic and obligatory buildup of streaming. For speech, this breakdown with large ISIs can occur even if speech tokens are spoken by one consistent talker (Best et al., 2008; Bressler et al., 2014; Lim et al., 2019), and introducing temporal gaps in speech reduces the facilitatory effect of talker continuity (Choi and Perrachione, 2019). Our evidence suggests that any discontinuities in the acoustic stream, whether due to temporal or featural aspects of incoming sounds, trigger involuntary reorienting of attentional focus, consistent with a domain-general stimulus-driven attention and auditory streaming account of talker-specific speech processing.

In addition, we found that the pupil dilation response during speech encoding was affected by talker discontinuity. Especially when listeners had to rapidly process incoming speech (i.e., for stimuli with the 0-ms ISI), we found a significant increase in pupil dilation when listeners were encoding speech spoken by mixed talkers compared to when it was from one consistent talker (**Figure 6**). This evidence provides further support for the idea that exogenous attentional reorientation is a critical cognitive mechanism engaged in processing speech in mixed-talker contexts. Furthermore, given the robust link between pupil dilation and activity in the LC-NE system (Aston-Jones and Cohen, 2005; Joshi et al., 2016; Costa and Rudebeck, 2016), our results suggest that rapid processing of mixed-talkers’ speech is mediated by the pupil-linked LC-NE response.

Our experiment provides the first psychophysiological evidence that a specific, domain-general neural system for attention and arousal influences cognitive operations during processing of talker variability in speech. One of the functional roles of the pupil-linked LC-NE system is to maintain and update the perceptual model of the surrounding environment (Bouret and Sara, 2005; Dayan and Yu, 2006; Sara and Bouret, 2012). The pupil dilation response serves as an index for NE release within the LC (Aston-Jones and Cohen, 2005; Samuels and Szabadi, 2008; Gilzenrat et al., 2010). NE is the neuromodulator known to serve as an interruption signal that halts ongoing top-down processes in the brain; this in turn, allows the system to automatically and exogenously switch attention to abrupt changes in sensory surroundings (Dayan and Yu, 2006; Sara and Bouret, 2012; Bouret and Sara, 2005). Consistent with this account, pupil dilation responses not only reflect the degree to which salient and surprising sound events draw attention involuntarily (Liao et al., 2016a; Huang and Elhilali, 2017), but also track rapid and unexpected changes in the auditory environmental structure even when the changes are behaviorally irrelevant (Zhao et al., 2019). Our results suggest a novel, but consistent perspective on parsing speech with talker discontinuity (and hence talker variability): mixed talkers’ speech appears to involuntarily reorient listeners’ attention the acoustic features of an unexpected, novel speech source (i.e., talker), and this process is, in part, accomplished by the pupil-linked LC-NE system.

### Consideration of other potential explanations for the cost of processing mixed talkers’ speech

Among previous psycholinguistic models that have been proposed to explain how talker variability is accommodated during speech processing, each posits a different cognitive mechanism to account for the additional processing cost of mixed-talker speech. These mechanistic explanations include uncertainty in accessing long-term, talker-specific memory representations (Kleinschmidt and Jaeger, 2015), increased computational complexity in resolving acoustic-phonetic ambiguity (Nearey, 1989; Johnson 2005), and higher demand for working memory resources to maintain multiple, potential acoustic-phonetic interpretations of upcoming speech (Nusbaum and Magnuson, 1997). However, in this study we did not see clear evidence of the sorts of neurophysiological signatures one would expect to be associated with the cognitive mechanisms posited by these models in either the ERP or neural oscillatory changes measured in response to talker variability.

The classical model describing how listeners resolve acoustic-phonetic ambiguities introduced by cross-talker differences is called *talker normalization* (e.g., Johnson, 2005; Nusbaum and Magnuson, 1997; Pisoni, 1997). This framework posits that listeners actively utilize intrinsic information (i.e., acoustic patterns) of speech signal (Ainsworth, 1975; Nearey 1989; Syrdal and Gopal, 1986) and extrinsic information from preceding speech context (e.g., Johnson 2005; Sjerps et al., 2013) to configure a talker-specific mapping between acoustics and phonemic representations. Thus, processing mixed-talker speech would lead to higher computational demands in order to continuously adjust and “normalize” the phonemic representations of each successive talker based on their unique acoustic features (i.e., speech formants; Sussman, 1986). According to this explanation, the computational cost of processing differences in mixed- vs. single-talker’s speech should likely manifest in early auditory processing.

One neural signature likely relevant to talker normalization is the magnitude of the N1 component of the auditory ERP, arising around ~100 ms after a sound onset. Extensive research has demonstrated that the auditory N1 response, associated with basic auditory perception (Hillyard & Picton, 1978; Näätänen & Picton, 1987; Pantev et al., 1996; Hyde, 1997; Martin et al., 2008), is modulated by top-down attention (Hillyard et al., 1973; 1998; Choi et al., 2013; 2014; Woldorff et al., 1993), and reflects the extent of neural adaptation in the auditory cortex (Maess et al, 2007; Pantev et al., 1988). As the N1 response magnitude is sensitive to changes in the acoustic features (repeating vs. non-repeating sound events; Todorovic et al., 2011; Herrmann et al., 2015), we would expect that the computational demands of forming talker-specific acoustic-phonetic representations in mixed- vs. single-talker’s speech would be reflected in differences in N1 magnitude. However, we did not find any such differences between the two talker conditions (**Figure 3**).

Although we did not observe any difference in the N1 magnitude between the talker conditions, it is worth noting some considerations regarding the N1 response, as the existing evidence is somewhat mixed. Previous EEG studies in the context of talker discontinuity (hence, variability) in speech have observed larger N1 magnitudes when parsing mixed-talker compared to single-talker speech (e.g., Kaganovich et al., 2006; Uddin et al., 2018; Zhang et al., 2013), or when listeners encountered an unexpected change in the talker of an attended speech stream (Mehraei et al., 2018). In contrast, other work has reported the opposite pattern: processing mixed talker speech either reduced (Shao and Zhang, 2019) or did not affect N1 magnitude (Zhang et al., 2013). One potential source of this discrepancy might arise from the amount of acoustic variability in the stimulus set, as the early evoked neural responses up to ~200 ms post-stimulus are contingent on the characteristics of the stimulus (Näätänen and Picton, 1987; Digeser et al., 2009; Khalighinejad et al., 2017; Trembley et al., 2003). Given the many factors that affect the N1, future studies are necessary to disambiguate the unique contributions of acoustic variability across and within talkers to markers of early auditory processing like the N1—as well as to speech processing efficiency.

Our findings on how talker discontinuity influences neural alpha oscillatory power do not seem consistent with the cognitive mechanism proposed by the other prominent model accounting for resolving talker differences in speech, that processing talker variability in speech is handled by an *active control process* (Nusbaum and Nusbaum, 1997). This model posits that listeners pre-allocate cognitive resources in order to flexibly resolve potential acoustic-phonemic ambiguity in speech signals, and to maintain robust speech perception accuracy in the face of variation (Nusbaum and Magnuson, 1997; Magnuson and Nusbaum, 2007; Heald et al., 2014). Thus, when listeners encounter mixed talkers’ speech, there is greater demand on working memory as listeners must simultaneously entertain multiple interpretations of incoming speech simultaneously in working memory. From this account, processing mixed talkers’ speech should presumably enhance alpha oscillatory power during speech encoding, as increasing working memory demand leads to parametric increases in alpha power (Jensen et al., 2002; Tuladhar et al., 2007; Obleser et al., 2012). However, we did not find any alpha power differences between the single and mixed talker conditions when listeners encoded the speech. Instead, we found the opposite pattern—a decrease in alpha power for mixed-compared to single-talker speech— during speech memory retention (**Figure 5**). This finding is inconsistent with increased working memory load as a mechanism for accommodating talker variability as suggested by the active control framework, but is consistent with the attentional enhancement of working memory maintenance (Lim et al., 2015; 2018).

One potential interpretation of our finding that alpha power during retention is lower for mixed- vs. single-talker speech sequences is that talker discontinuity disrupts attention directed to working memory. Attention enables effective encoding and maintenance of relevant information in working memory (Awh and Jonides, 2001; Serences and Kastner, 2014; Gazzaley and Nobre, 2012), whereas attentional disruption can impair working memory maintenance. Also, attention directed to working memory items improves memory recall of stored items (Oberauer and Hein, 2012; Lim et al., 2018; 2021), and the degree of attentional benefit on memory recall can be related to alpha oscillatory power enhancement (Lim et al., 2015). As auditory-based attention depends on the ability to form a coherent auditory object (Shinn-Cunningham, 2008), it is possible that emergence of a single auditory stream via talker- and temporal-continuity might also facilitate efficient storage of the resulting coherent object in memory. In contrast, talker and/or temporal discontinuities that break down streaming may impair efficient storage and allocation of attention to working memory objects.

Although our present results are not consistent with the predictions of the active control framework (Nusbaum & Magnuson, 1997), it is worth noting that this framework may not be mutually exclusive from the auditory streaming account. Recent behavioral research demonstrates that both auditory streaming and active control processes appear to work in parallel to support processing of talker variability in speech (Kapadia & Perrachione, 2020; Choi, Kou, & Perrachione, submitted). For instance, when listeners must parse speech in the presence of background noise or when they have an ongoing expectation of uncertainty about the upcoming talker, they seem to pre-allocate cognitive resources to cope with the ongoing listening challenge. This enhanced cognitive load can tie up available resources, which can then impede the attentional benefit that listeners otherwise gain when processing speech that is continuous in talker (Kapadia & Perrachione, 2020).

Finally, the *ideal adapter framework* (Kleinschmidt & Jaeger, 2015) formalizes the longstanding episodic approach to understanding processing variability in speech perception (e.g., Goldinger, 1996, 1998). These approaches assert that listeners maintain long-term memory representations of talkers’ speech, against which the incoming speech signal is compared to support recognition. According to this framework, the additional processing cost of mixed-talker speech is associated with the greater number of possible competing interpretations (or “models”) of a given speech token. By anticipating the speech of a particular talker (or class of talker) in advance, listeners can reduce the decision space to a subset of these (e.g., talker-specific) models, leading to more efficient speech processing. However, the current formulation of this framework does not address how its model-selection operations might be implemented in specific neurobiological processes that are distinct from the input or processing predictions made by talker normalization or the active control hypotheses, as discussed above.

## Conclusions

The present work shows that talker discontinuity in speech interferes with both immediate processing of, and subsequent working memory for, speech. Differences in ERP and pupil dilation responses suggest that the behavioral costs associated with processing variable, mixed-talker speech are the result of added auditory processing demands incurred by automatic attentional reorientation to the new source of speech upon talker discontinuity. Neural alpha oscillatory power results suggest that the interference effect of talker variability in speech is also present when listeners maintain speech information in working memory after speech encoding. Collectively, our results demonstrate that talker changes evoke an involuntary reorientation of attention via domain-general processes, which interact with the pupil-linked LC-NE system in determining processing efficiency of mixed-talker vs. single-talker speech.

## Conflict of Interest

The authors declare no competing financial interests.

## Acknowledgement

This work was supported by NIH grants (R03DC014045, R01DC004545) and Brain and Behavioral Research Foundation NARSAD Young Investigator grant to TKP, and NIH grant (R01DC009477) to BGSC. SJL was supported by grants from the NIH (T32DC013017) and the NSF (BCS 1840674).

## Notes

### Competing Interest Statement

The authors have declared no competing interest.

