## Supplementary Material for "Talker discontinuity disrupts attention to speech: Evidence from EEG and pupillometry"

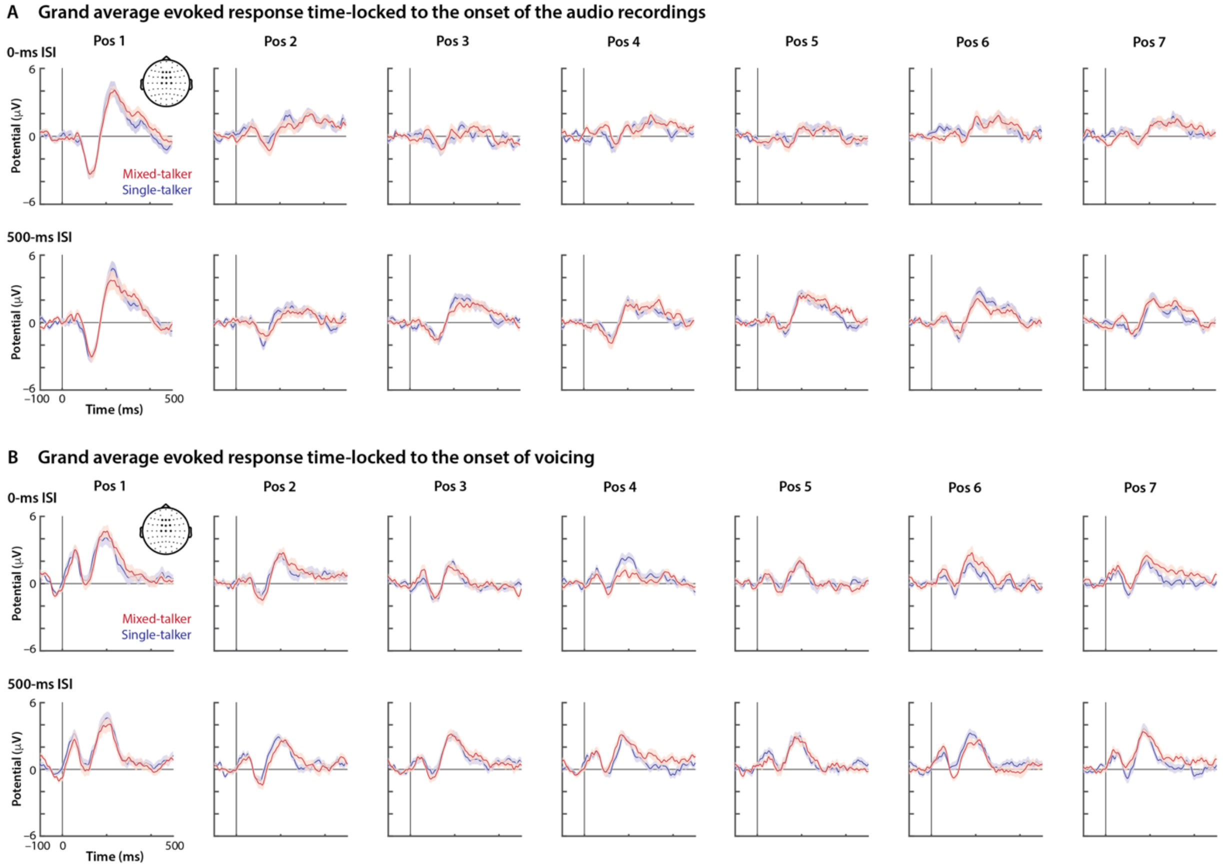


**Figure S1.** Grand average evoked responses to spoken digits across the sequence positions (position 1–7) in the 2 *talker* × 2 *temporal discontinuity* conditions. The evoked responses are computed based on the neural responses time-locked to the onset of the audio recordings (A) and time-locked to the onset of voicing of the stimulus recording. Note the differences in the evoked response patterns and the response amplitudes of the first spoken digit compared to those of the rest of the digits in the sequence (position 2–7). The evoked responses were extracted from nine fronto-centro scalp channels highlighted in the topographical map. The error bars indicate ±1 SEM.

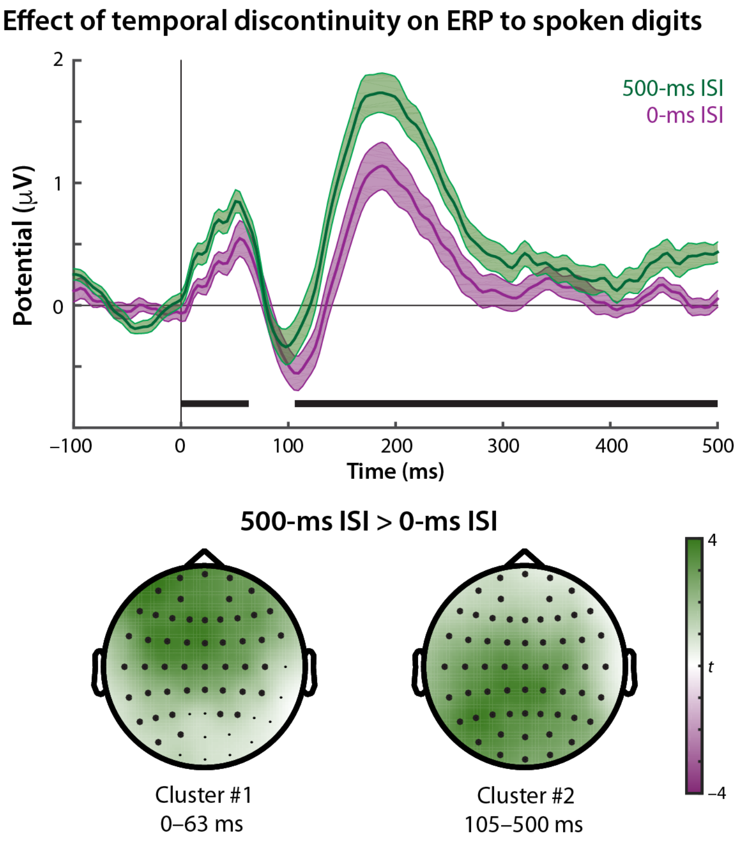


**Figure S2.** The main effect of temporal discontinuity on the evoked response to spoken digits in the sequence (time-locked to voicing onset of speech). Top, the grand average time courses in the 500-ms vs. 0-ms ISI conditions. Error bars indicate ±1 SEM. Bottom, the topographical maps of the two clusters exhibiting a significant contrast between the 500-ms ISI vs. 0-ms ISI conditions. The distributions of *t*-statistics of the contrast are illustrated.

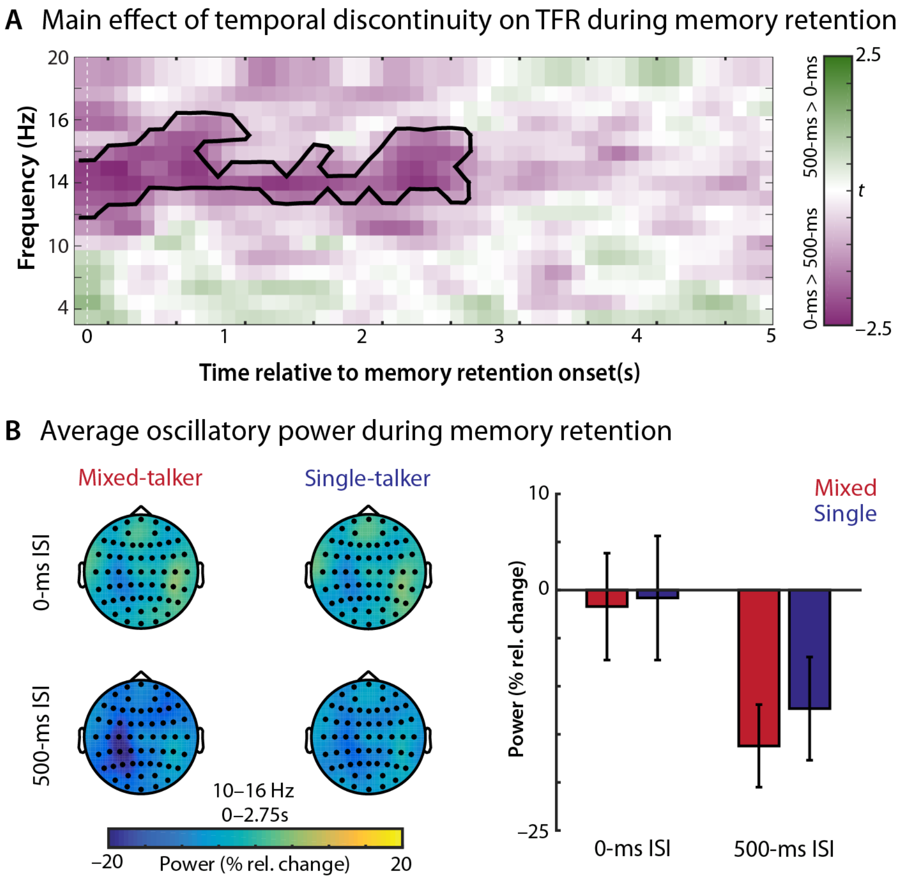


**Figure S3.** The main effect of temporal discontinuity on the neural oscillatory power during the 5-s memory retention phase. **(A)** The time–frequency representation of the cluster exhibiting a significant 0-ms > 500-ms ISI contrast (aggregated across the two talker conditions) during memory retention. The colors indicate the distributions of *t*-statistics of the 500-ms vs. 0-ms ISI contrast. **(B)** Average oscillatory power (10–16 Hz around 0–2.75s with respect to the memory retention phase) in each of the 2 *talker* × 2 *temporal discontinuity* conditions over the significant 0-ms ISI > 500-ms ISI cluster shown in (A). Left, topographical illustrations of the condition-specific averages of oscillatory power over the frequencies and times belong to the cluster. Right, a bar plot of condition-specific oscillatory power average over the electrodes, frequencies, and times of the cluster. Error bars indicate ±1 SEM.
